# Multimodal Ganzfeld-induced visual experiences are associated with accompanying mental experiences and distinct EEG microstate dynamics

**DOI:** 10.64898/2026.08.28.747793

**Authors:** Xinlin Wang, Yannick Pomorin, Emma Peters, Daniel Erlacher, Thomas Koenig

## Abstract

During wakefulness, we are used to perceive the environment through our senses, act on it and take these inputs to update our experiences and build the perceptions. When the inputs are not longer accurate or structured, people would sometimes have hallucinatory experiences. Whether such experiences are associated with distinct patterns of thought, and how they relate to large-scale brain dynamics, remains unclear. To address these questions, we combined experience sampling protocol with EEG recording during multimodal Ganzfeld, where participants were exposed to unstructured, uniform visual and auditory stimulation. Participants repeatedly reported the complexity of their visual experiences together with ongoing thoughts related to perceptual belief, prediction–perception mismatch, active updating, and prior mentation. EEG microstates were extracted to characterize the temporal dynamics of large-scale brain networks. We found that visual complexity was related to some of these dimensions, but partly distinct in simple and complex visual experiences. These phenomenological changes were accompanied by distinct, and often nonlinear, dynamics of large-scale brain networks involved in visual processing, salience detection, and internally directed cognition. It also indicates that this paradigm might be a valuable model for investigating the mechanisms underlying hallucinatory experiences in psychosis.

## Introduction

The brain is increasingly understood as a self-organizing system whos ongoing activity is continuously constrained by the external world rather than initiated by it (Miskovic, Lynn, et al., 2019). Ample evidence indicates that spontaneous brain activity in the absence of external input also bears important functional significance (see review in Moutard et al., 2015). This activity can be characterized on many scales from single neurons to large-scale brain networks (Miskovic et al., 2019), in computational simulations (Dehaene & Changeux, 2005), and it is accompanied by subjective mental representations. These characterizations can be understood through a theoretical framework called predictive coding (K. Friston, 2010), according to which the brain continuously generates top-down predictions from an internal model, compares them against incoming sensory signals or representations, and propagates the resulting predictive errors upward through a hierarchy to update the internal model. Consequently, perception is considered an inferential process in which sensory evidence and prior expectations interact to estimate the most likely causes of experience (K. Friston, 2010). How we perceive the external world emerges from the balance between bottom-up and top-down stream processing.

When this balance is disrupted, it may cause abnormal perceptual experiences, such as hallucinations (K. J. Friston, 2005; Sterzer et al., 2018). Traditionally, the mechanisms underlying hallucinations have primarily been investigated in clinical populations, particularly in schizophrenia patients as a core symptom (Frith, 2005; Zmigrod et al., 2016). Evidence from schizophrenia or voice-hearing populations suggested that hallucinations were associated with alterations in predictive inference (Sterzer et al., 2018). The findings differred depending on whether abnormalities were primarily reflected by overly strong effects of top-down predictive signals on neural activity in sensory cortex (Horga et al., 2014; Shergill et al., 2014) or a failure to attenuate predictable internal signals in the somatosensory cortex (Ford & Mathalon, 2005; Hubl et al., 2007; Powers et al., 2017; Sterzer et al., 2018). Hallucinations (in schizophrenia) have also been linked to impaired sensory gating, reflecting reduced filtering of irrelevant sensory information (Javanbakht, 2006). Previous studies have largely examined hallucinations by manipulating prior expectations in specific experimental tasks, which made it difficult to disentangle the contribution of altered predictive inference from more general deficits in sensory processing. Besides, hallucinations in clinical populations are often accompanied by broader psychosis symptoms, making it challenging to determine whether effects are specifically associated with hallucinations or psychosis more broadly (Powers et al., 2017).

Sensory deprivation paradigms, such as multimodal Ganzfeld (MMGF), provide a safe opportunity for inducing hallucinatory experiences in healthy individuals by exposing them to an unstructured, uniform sensory environment (Miskovic, Lynn, et al., 2019; Wackermann et al., 2008). Typically elicited with homogeneous lights and sounds, the MMGF can induce a range of perceptual hallucinatory experiences, from colors and geometric patterns to more vivid, dream-like experiences (Lloyd et al., 2012; Wackermann et al., 2002). As discussed above, such conditions are thought to reduce the precision of sensory signals, resulting in a relatively flat likelihood distribution in which assumingly random prior expectations may be increasingly weighted during perceptual inference. MMGF therefore offers a valuable framework for investigating how internally generated activity contributes to hallucinatory experiences in the healthy brain when external sensory constraints are weakened.

Previous EEG studies have primarily reported changes in parietal and occipital alpha activity during MMGF, suggesting alterations in attentional and memory-related processes (Miskovic, Bagg, et al., 2019; Pütz et al., 2006). However, increasing evidence showed that cognitive processing, such as attention and memory, is not executed by a single brain region or single brain network, but rather by several largely non-overlapping brain networks (Marek & Dosenbach, 2018). From this perspective, local oscillatory changes alone might be insufficient to understand the neural mechanisms underlying Ganzfeld-induced hallucinatory experience, which might involve widely distributed brain regions and networks. Partly supporting this view, a recent fMRI study found that MMGF was associated with reduced functional connectivity between the thalamus and primary sensory cortices, alongside increased connectivity within the default mode network (Schmidt et al., 2020). These findings suggested a shift from externally driven sensory processing toward internally generated activity during MMGF. However, the observed effects were related to MMGF-induced altered state of consciousness in general, not directly linked to hallucinatory experience. It remains unclear whether and how large-scale brain network dynamics are associated with Ganzfeld-induced hallucinatory experience.

Besides, previous studies have primarily focused on perceptual phenomena induced by MMGF (Lloyd et al., 2012; Shenyan et al., 2024; see review by Miskovic, Lynn, et al., 2019). Considering that hallucinatory experiences do not occur in isolation but are embedded within an evolving stream of consciousness or thoughts, the patterns of ideation accompanied by or preceded by perceptual phenomena might also carry important information for explaining their formation. Yet little is known about the relationship between perceptual phenomena and ongoing patterns of thought during MMGF, or whether both are associated with changes in large-scale brain-network dynamics.

To address these questions, we adopted a neurophenomenological approach that combined experience sampling with EEG recordings during MMGF in a healthy population (Fig. 1a). Participants were repeatedly asked to report not only their perceptual experiences but also ratings of their ongoing mentation related to predictive inference experience at random intervals (Fig. 1c). Drawing on the Prediction-Related Experiences Questionnaire (O’Brien et al., 2024), which was developed to assess subjective experiences of prediction in everyday life, we measured the extent of participant’s perceptual belief, perception prediction mismatch, and active updating of the new perception during MMGF, as well as prior mentation that preceded the perceptual experiences (Fig. 1b). We hypothesized that increased complex perceptual phenomena would be associated with increased belief in the reality of the experience and greater active updating, reflecting attempts to integrate emerging perceptual content into a coherent interpretation. Furthermore, if MMGF-induced perceptual phenomena are shaped by altered ongoing predictive processes, we expected them to be related to perception prediction mismatch and preceding patterns of mentation.

**Figure 1.**
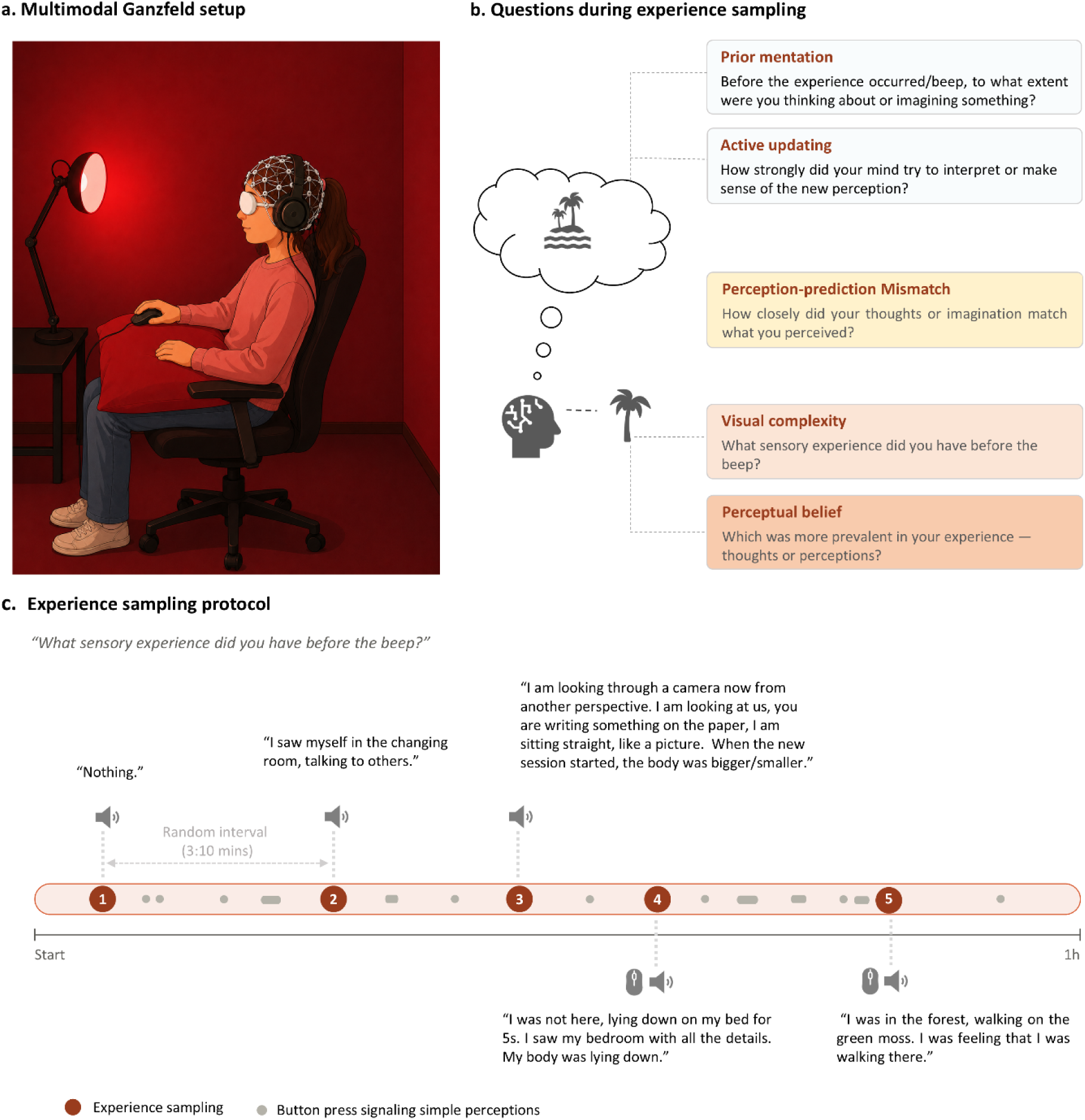
Experiment setting and study design. **a.** AI-generated illustration of Multimodal Ganzfeld setup. The participant was seated in a comfortable chair, wearing a 64-channel EEG cap, headphones delivering nature-raining white noise, and modified ping-pong ball halves covering the eyes. Participant indicated their perceptual experience during the session by a computer mouse held in the right hand. A pillow was placed on the participant’s lap to support their arms. **b.** Experience-sampling questions were used to assess the visual experience and characteristics of ongoing mental experience during the Ganzfeld session. The illustrated example shows a participant thinking about or imagining a natural beach and reporting perceptual experiences related to this mental content. This experience was measured using different questions: visual complexity, perceptual belief, perception-prediction mismatch, active updating, and prior mentation. **c.** Experience sampling protocol. It shows an example timeline from one participant during the 1-hour Ganzfeld session. The protocol was paused at pseudo-random intervals by a beep. Then the participant was asked to report their sensory experiences immediately before the beep and also answer other questions. During the session, participants were instructed to signal the appearance of complex visual experiences such as objects and scenes by right-clicking the mouse. This also triggered an experience sampling. They were also instructed to report simple visual experiences (e.g., colors, geometric shapes) by pressing and holding the left mouse button for the duration of each experience. Signaling simple percepts did not automatically trigger experience sampling. The orange circle indicates protocol interruptions with experience sampling reports, and the grey marks indicate the occurrence of simple visual experiences. Self-reports from one participant are shown above and below the timeline.

To identify the dynamics of large-scale brain networks associated with these phenomena, we conducted EEG microstate analysis on segments corresponding to the subjective experience reports. EEG microstates assumingly capture synchronized global networks and their dynamics over time by characterizing quasi-stable global patterns of scalp electrical activity that typically persist for approximately 60–120 ms before transitioning to another state (Michel & Koenig, 2018). EEG microstates have been linked to the coordinated activity of large-scale functional brain networks identified in fMRI studies (Custo et al., 2017; Michel & Koenig, 2018). It allows the study not only of the functional brain network but also of its temporal properties, such as duration, occurrence, and coverage. It has emerged as a promising tool for measuring the spontaneous fluctuations of activity in large-scale brain networks underlying different conscious experiences (Bréchet & Michel, 2022). Patients with schizophrenia who experienced hallucinations showed a particular signature in distinct EEG microstates. For example, a shorter duration of Microstate D in periods with hallucinations indicated a facilitation of the misattribution of self-generated inner speech to external sources during hallucinations (Kindler et al., 2011). Increased contribution of microstate C has been associated with higher hallucination scores in adolescents with 22q11.2 syndrome (Tomescu et al., 2014). Given that MMGF-induced visual experiences involve attentional, memory-related, and higher-order cognitive processes in addition to perceptual processing (Lloyd et al., 2012; Miskovic et al., 2019), we expected visual complexity to be associated with alterations in microstates linked to higher-order cognitive networks, particularly in Microstates C and D. Considering that simple visual experience during MMGF might be particularly associated with visual perception and partly originate from entoptic processes related to the visual system, whereas more complex and structured visual experience might recruit additional cortical processes (Miskovic et al., 2019), we hypothesized a non-linear relationship between visual complexity and the dynamics of microstate B, which has been associated with visual network activity. The neural correlates of perceptual belief, surprise, active updating, and prior mentation were considered exploratory.

## Results

### Complexity of MMGF-induced visual experiences was associated with ongoing mental experiences

We extracted 304 experience sampling trials from 54 participants. To investigate the relationship between ongoing mental experiences and the complexity of visual experiences observed during MMGF, we first rated visual experience complexity from participants’ self-reports during the protocol session using an adapted version of Lloyd et al.’s (2012) framework (Fig. 2). These experiences includes a continuum of visual perceptions, from abstract simple perceptions: 1) simple visual changes, e.g., color changes, flash; 2) Simple shapes or patterns e.g., lines, dots, cloudy and 3) Combine 1 and 2, e.g., a sequence of dots moving like a river; to external object related and immersive complex perceptions: 4) objects, e.g., face, tree, ghost and 5) environment, e.g., sea, garden or dream liked experiences. Trials in which participants reported no sensory experience were assigned a baseline score of 0 (Fig. 2).

**Figure 2.**
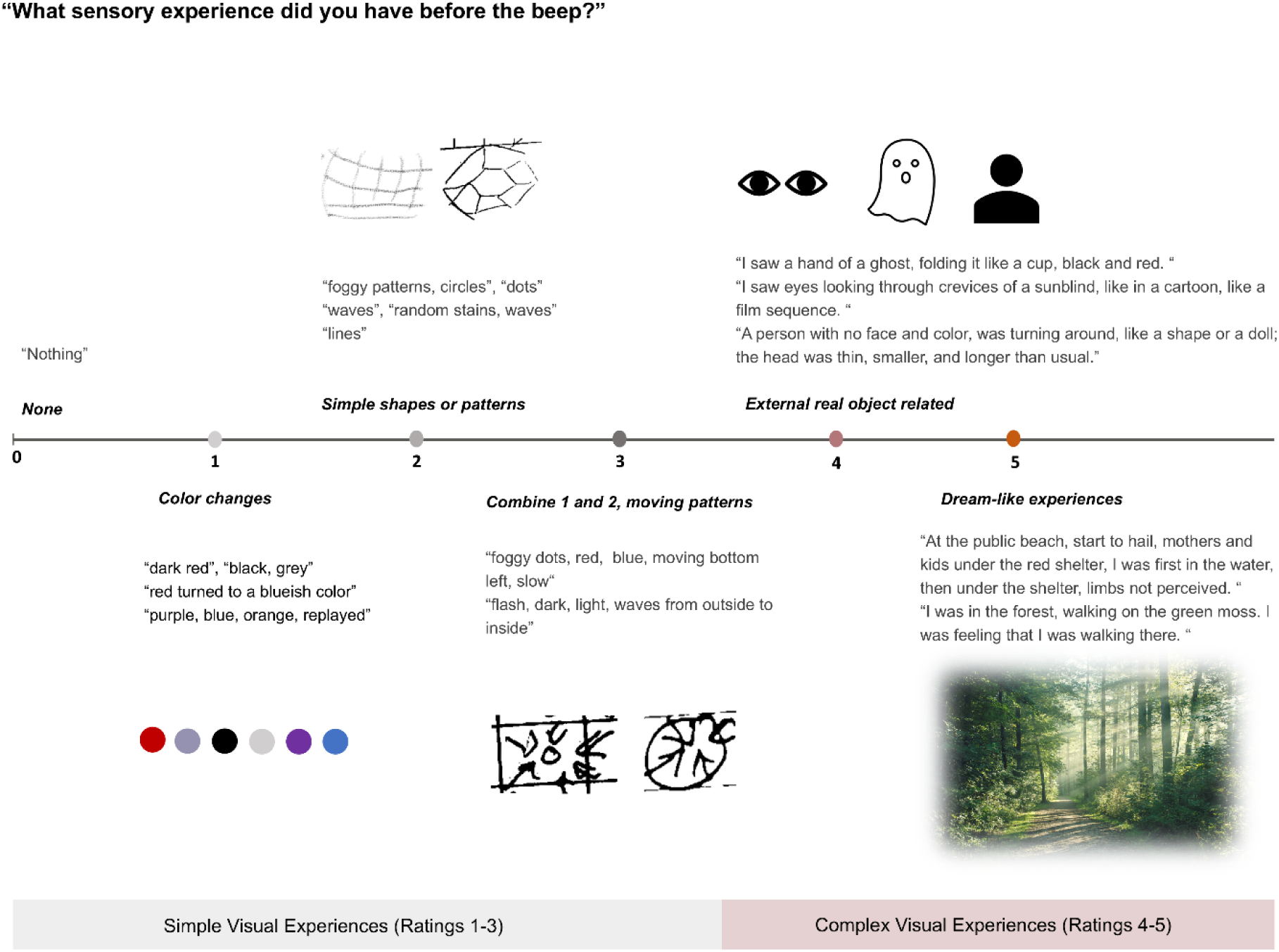
Visual experience complexity analysis from participants’ self-reports during MMGF. Illustration of each rating of Visual complexity, example episodes and figures from participants’ self-reports. The level of visual complexity was rated by researchers.

Because it remains unclear whether visual complexity has linear relationships with other ratings of ongoing mental experiences, we then fitted separate linear mixed-effects models with collected ratings as the dependent variables and visual complexity terms as the predictor. The linear and quadratic polynomial terms were used as fixed factors, with subject as a random factor.

First, we observed that the association between Perceptual belief and visual complexity was significant in the linear term (F(1, 290.3) = 63.29, p < .001), whereas the quadratic term was not significant (F(1, 301) = 1.01, p = .29). The estimated linear trend was positive (β = 9.37, t(289) = 7.89, p < .001). As shown in the figure (Fig. 3a), the result suggested that as visual experiences became more complex, participants increasingly tended to believe that their experiences were perceptions.

**Figure 3.**
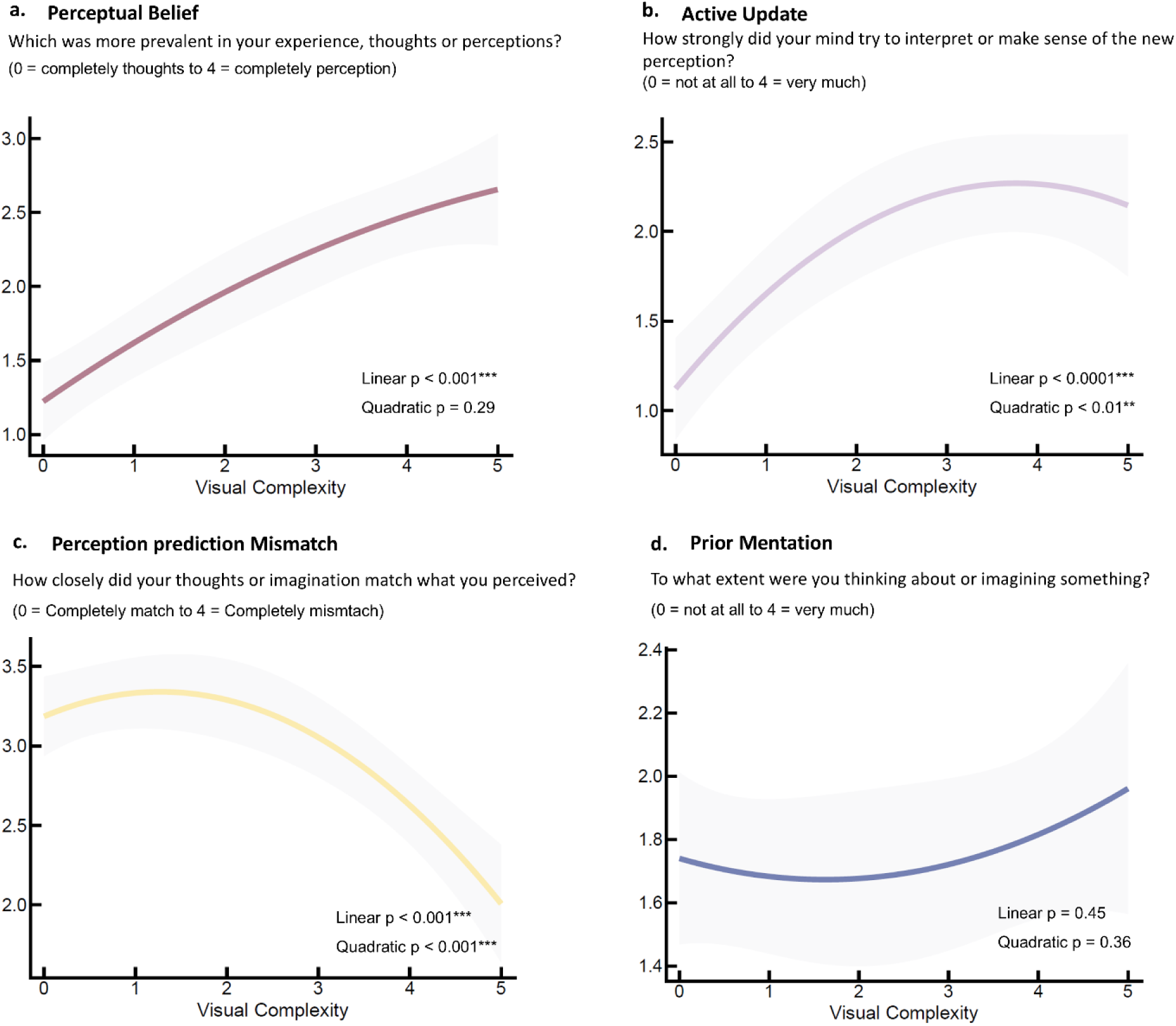
Relationship between visual complexity and ongoing mental experiences. **a–d** showed associations between visual complexity and the patterns of mentation: perceptual belief (**a**), active updating (**b**), perception–prediction mismatch (**c**), and prior mentation (**d**).

Then, we found that Active updating significantly increased as the visual experience became more complex in both the linear term (F_(1, 298.60)_ = 41.38, p < .001) and the quadratic term (F_(1, 299.18)_ = 9.16, p < .01). The estimated linear trend was positive (β = 7.76, t_(299)_ = 6.38, p < .0001), and the quadratic trend was negative (β = −3.37, t_(299)_ = −3.00, p < .01). The results suggested that participants increasingly interpreted and made sense of their new perceptual experiences as their visual experiences became more complex overall, but that this increase peaked when complex experiences began to emerge (Fig. 3b).

For the Perception and prediction Mismatch, we observed significant effects in both linear (F_(1, 277.74)_ = 23.12, p < .001) and quadratic (F_(1, 299.68)_ = 13.24, p < .001) terms, with the significant negative linear trend (β = −5.62, t_(277)_ = −4.76, p < 0.0001) and negative quadratic trend (β = −3.98, t_(300)_ = −3.61, p < .001). The results showed that participants reported a relatively stable mismatch between their expectations and perceptual experiences during simple visual phenomena, such as color changes and geometric patterns. At higher levels of visual complexity, mismatch decreased with visual complexity, indicating that complex visual experiences were increasingly aligned with participants’ ongoing thoughts and expectations (Fig. 3c).

For the Prior mentation, we did not find a significant linear association with visual complexity (F(1, 286.82) = 0.56, p = .45) or a quadratic association (*F*(1, 300.87) = 0.83, p = .36). It suggested that MMGF-induced visual experiences were not affected by prior mentation.

### Seven class microstates captured global network changes among the experience sampling

For each experience-sampling trial, we extracted the EEG segment corresponding to the 30 s preceding each beep probe for microstate analysis. We then clustered microstate template maps from 4- to 7-class solutions across participants and experience-sampling trials (Fig. S1). We had an explicit hypothesis about microstates B, C, and D and visual experience during MMGF. Only the seven-class solution included all the microstate class maps (microstates A, B, C, D, E, F, G) we hypothesized; we therefore chose the seven-class solution (Fig. 4a) for further analysis.

**Figure 4.**
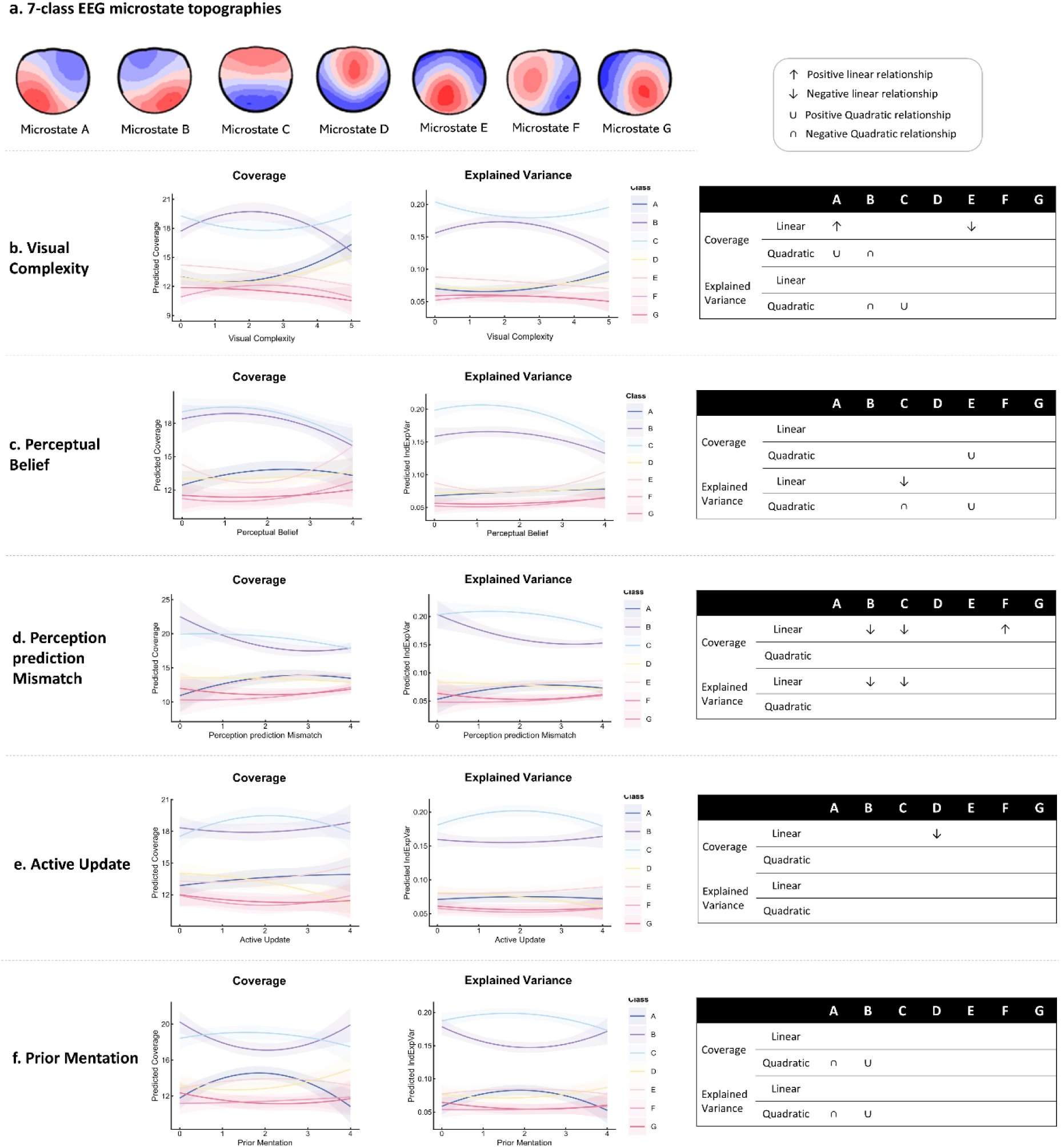
Results of microstate analysis. **a.** The seven EEG microstate classes identified by k-means cluster analysis across participants and the two temporal microstate parameters used for subsequent analyses: coverage (%) and explained variance. **b–f** showed associations between microstate parameters and experience sampling ratings: visual complexity (**b**), perceptual belief (**c**), perception–prediction mismatch (**d**), active updating (**e**), and prior mentation (**f**). Coloured lines represent microstate parameter values for each microstate class, and shaded regions indicate 95% confidence intervals. Summary tables on the left indicate significant linear and quadratic associations identified by mixed-effects models after FDR correction. Positive and negative linear effects are denoted by upward and downward arrows, respectively, whereas positive and negative quadratic effects indicate U-shaped and inverted U-shaped relationships.

### Visual complexity and ongoing mental experiences were associated with distinct linear and nonlinear changes in microstate dynamics

To examine large-scale brain network activity associated with visual complexity during MMGF, the average coverage of the EEG signal in percent (Coverage) and the explained variance for each microstate class (Explained Variance) were extracted. We conducted separate linear mixed-effects models for each experience-sampling rating. The initial visualization of the data suggested that the relationships between experience-sampling ratings and microstate measures were not uniformly linear. To assess potential associations between distinct microstate networks and experience-sampling ratings, we decomposed the ratings into linear and quadratic (poly, degree = 2) terms. The results are depicted in Fig. 4b-f.

#### Visual complexity

Firstly, the mixed-effects models revealed a significant interaction between microstate classes and the linear term on coverage (F_(6, 2107)_ = 4.13, p < .001). The post hoc test showed a significant positive linear effect on coverage of microstate A (β = 41.01, t_(1875)_ = 3.18, p = .010) and a negative effect on microstate E (β = -32.87, t_(1875)_ = -2.55, p = .037). Then, there was also a significant interaction between microstate classes and the quadratic term (F_(6, 2107)_ = 5.77, p < .001). Post-hoc tests revealed that microstate B showed a significant negative quadratic effect (β = −52.71, t_(2010)_ = −4.09, p = .0003), suggesting that Coverage in microstate B was highest at intermediate levels of Visual Complexity and decreased toward both the lower and higher extremes (an inverted U-shape). Microstate A showed a significant positive quadratic effect (β = 32.25, t_(2010)_ = 2.50, p = .043), suggesting that Coverage in microstate A was lowest at intermediate levels of Visual Complexity and increased toward both extremes (U-shape). For explained variance, the mixed effects model revealed a significant interaction between microstate classes and the linear term (F_(6, 2067)_ = 3.40, p = .002). However, there was no significant linear effect on the specific microstate class after the corrected post hoc test. Then, the model also showed a significant interaction between microstate classes and the quadratic term (F_(6, 2067)_ = 4.97, p < .001). The following tests revealed a significant negative quadratic effect for microstate B (β = −0.54, t_(2031)_ = −4.01, p = .0004), and a significant positive quadratic effect for microstate C (β = 0.35, t_(2031)_ = 2.62, p = .031) (Fig. 4b).

#### Perceptual Belief

Firstly, the mixed-effects model revealed a significant interaction between microstate class and the linear term for coverage (F_(6, 2107)_ = 2.81, p = .010), but there was no significant linear effect for specific microstate classes in the post hoc test. Then the model showed a significant interaction between microstate classes and the quadratic term (F_(6, 2107)_ = 3.28, p = .003), and the following post hoc test revealed a significant positive quadratic effect on microstate E (β = 40.26, t_(2089)_ = 3.11, p = .013), where the coverage of microstate E was lowest at the intermediate levels of Perceptual belief. For explained variance, the mixed effects model revealed a significant interaction between microstate classes and the linear term (F_(6, 2066)_ = 4.89, p < .001), and the quadratic term (F_(6, 2066)_ = 3.96, p = .001). Post hoc test showed both a significant negative linear effect (β = −0.59, t_(2009)_ = −4.36, p < .001) and negative quadratic effect (β = −0.45, t_(2103)_ = −3.32, p = .006) for microstate C, and a significant positive quadratic effect for microstate E (β = 0.36, t_(2103)_ = 2.64, p = .029) (Fig. 4c).

#### Perception prediction Mismatch

The mixed effects model revealed a significant interaction between microstate classes and the linear term (F(6, 2107) = 4.79, p < .001), and the quadratic term (F(6, 2107) = 2.27, p = .034) for coverage. The following tests showed significant negative linear effects of microstate B (β = −35.77, *t*_(2016)_ = −2.77, p = .020) and microstate C (β = −36.31, *t*_(2016)_ = −2.81, p = .020), and a significant positive effect of microstate F (β = 30.80, *t*_(2016)_ = 2.38, p = .040). However, we did not observe the significant quadratic effects on specific microstate classes after the post hoc test. For explained variance, the mixed-effects model showed a significant interaction between microstate class and the linear term (F_(6, 2067)_ = 5.06, p < .001). The post hoc test revealed a significant negative linear effect of microstate B (β = -0.47, *t*_(2041)_ = −3.30, p = .003) and C (β = -0.49, *t*_(2041)_ = −3.61, p = .002) (Fig. 4d).

#### Active update

The mixed effects model only reveal a significant effect between microstate class and the linear term of coverage (F_(6, 2107)_ = 2.41, p = .025). Post hoc analysis showed a significant negative linear relationship between coverage of microstate D and active updating (β = -37.37, *t*_(1846)_ = -2.877, p = .029). For explained variance, the mixed-effects model did not show a significant interaction between microstate class and the linear term (F(6, 2067) = 1.00, p = .42) or the quadratic term (F_(6, 2067)_ = 1.62, p = .14) (Fig. 4e).

#### Prior mentation

For coverage, the mixed-effects model showed only a significant interaction between microstate class and the quadratic term (F_(6, 2107)_ = 6.12, p < .001). Post hoc tests revealed a significant negative quadratic effect for microstate A (β = −52.01, t_(1975)_ = −4.02, p = .0004) and a positive quadratic effect for microstate B (β = 47.80, t_(1975)_ = 3.70, p = .0008). Similarly, for explained variance, the model only revealed a significant interaction between microstate class and the quadratic term (*F*_(6, 2067)_ = 4.90, p < .001). The post hoc test also showed a significant negative quadratic effect for microstate A (β = −0.44, t_(2003)_ = −3.21, p = .005) and a positive quadratic effect for microstate B (β = 0.45, t_(2003)_ = 3.29, p = .005) (Fig. 4f).

### Functional characterization of microstate classes based on topographic similarity

To associate cognitive functions more objectively between different microstates, we performed a topographic similarity analysis between our class microstates and a database of published microstate maps (Koenig et al., 2023). We then extracted the empirical findings associated with the published maps showing topographic similarity to our microstate maps. Findings from studies involving cognitive tasks, schizophrenia, and altered states of consciousness were selected. The results (shown in Supplementary Tables 1-7) serve as a reference for functional interpretation of each microstate class observed in the present study. We ordered studies by descending similarity to our template maps, and weighted findings from higher-similarity studies more heavily during interpretation, with threshold of 80% similarity. Because the number of available studies and the range of similarity differ across microstates, we compared some template microstates with fewer published topographies, which allowed a lower threshold of 70%. Empirical findings from studies with higher similarity to our template suggested that microstate A is associated with temporal cortex-related processing, microstate B with primary visual processing, microstate C with task-positive and salience processing, microstate D with cognitive control, microstate E with emotional, arousal, and salience-related processing, microstate F with default mode network-related processing, and microstate G with somatosensory network. The appearance of microstates seems to be modulated differently during eyes open and eyes closed conditions (Seitzman et al., 2017). Because EEG in the present study was recorded with eyes open, comparisons with studies conducted under eyes clossed conditions should be interpreted with caution (See Eyes column in Supplementary Tables 1–7).

## Discussion

The present study combined a neurophenomenological approach with EEG microstate analysis to investigate the phenomenological and EEG microstate correlates underlying MMGF-induced visual experiences in healthy individuals. At the phenomenological level, in general, greater visual complexity was accompanied by higher levels of active updating, increased perceptual belief, and reduced prediction–perception mismatch. But these relationships differed slightly between simple and complex visual experiences, indicating that MMGF-induced simple and complex visual experiences might recruit partly distinct cognitive processes. This idea was confirmed by the findings from the EEG microstate analysis, where visual experience complexity was characterized by dynamic changes in microstates A, B, C, and E. We also observed distinct associations of microstates with perceptual belief, prediction–perception mismatch, active updating, and prior mentation.

From the Bayesian inference perspective, hallucinations can be modeled as strong priors that occur in the absence of sufficient sensory evidence (K. J. Friston, 2005). Recent studies also suggested that hallucinations might be associated with false beliefs about the precision of priors (Yon & Frith, 2021; Powers et al., 2017). And precision could be modulated by contextual factors such as attention (see review in Auksztulewicz & Friston, 2016). In the present study, participants were exposed to a highly ambiguous and imprecise sensory environment in which sensory evidence was absent. Participants might thus have increasingly relied on internally generated predictions to build their experiences. Increased attention to their experience and efforts to make sense of the ambiguous perceptions might have further strengthened these predictions and increased the subjects’ reliance on them. In the current study, we observed that increased complexity of visual hallucinations was associated with stronger active updating, suggesting that increased efforts to interpret ambiguous perceptions might contribute to the emergence of complex visual experiences during the MMGF. But it is also worth noting that we did not directly measure the precision or perceived reliability of predictions in the current study. Future studies could test this hypothesis more directly by a dedicated study design or by incorporating computational models to quantify prediction precision during MMGF.

Besides, we observed a negative relationship between visual complexity and Perception-prediction mismatch: When visual complexity increased, participants reported progressively lower mismatch between their thoughts or imagination and what they perceived. This relationship was mainly observed in complex visual experiences. One possible explanation might be that more complex visual experiences emerge when the brain’s generative model has formed a relatively stable representation, such that prediction errors are fading out. However, under the MMFG settings, highly imprecise sensory inputs led to constantly imprecise prediction errors that remained unreliable for updating the internal model (Auksztulewicz & Friston, 2016). Thus, the formation of complex visual experiences in MMFG are unlikely to be explainable terms of reduced prediction errors. Also, it remains unclear whether a close match between prediction and perception at the level of subjective phenomena in the present study implies fewer prediction errors at the neural level. An alternative explanation of the negative relationship might be that more internal and effortful attention to increasingly complex visual experiences resulted in reduced monitoring of the correspondence between the prediction and perception.

Another factor that might explain participants’ hallucinational experience is the failures in source monitoring or reality monitoring (Brookwell et al., 2013; Garrett & Silva, 2003). Source monitoring refers to a set of meta-cognitive processes used to make inferences about the origin of internally and externally generated events (Johnson et al., 1981, 1993). According to this account, participants prone to hallucination are not only impaired in their capacity to discriminate between internally and externally generated events, but also present a specific cognitive bias towards the misattribution of internally generated experiences to external sources (Bentall, 1990). In our results, we observed that increased visual complexity was associated with stronger perceptual belief, where participants increasingly perceived their experience as external perception, not internally generated thoughts. Besides, as visual complexity increased, participants reported a closer match between their thoughts or imagination and what they perceived, suggesting a progressive blurring of the external/internal boundaries.

Increased activation of sensory areas during hallucination experiences has been frequently reported (Allen et al., 2008; Dierks et al., 1999; Horga et al., 2014). However, the present study observed a non-linear relationship between the complexity of visual hallucinations and the coverage of Microstate B. This microstate class has been previously attributed to the visual network, specifically to early visual areas (Custo et al., 2017). The coverage and explained variance of microstate B as function of the complexity of the hallucinations followed an inverted-U function and reached its maximum at moderate levels of visual complexity (simple hallucination). These results fit the hypothesis that simple hallucinations manifested as lights, colors, or geometric designs initiated in the occipital regions (early visual areas) (Degueure et al., n.d.; ffytche, 2008), and therefore may be associated with increased appearance of microstate B. Complex hallucinations presented as formed images of objects, animals, people, or dream-like scenes, are instead rarely occipital in origin and generally arise from higher order perceptual regions, such as anterior ventral temporal regions (ffytche, 2008). Also, complex visual hallucinations in Charles Bonnet syndrome have been associated with increased activity in the lateral temporal cortex, striatum and thalamus (Adachi et al., 2000), and patients with temporal lobe epilepsy may also report complex visual hallucinations (Nelson et al., 2016; Revdal et al., 2020). Therefore, in the present study, the reduced coverage and explained variance of microstate B observed at higher levels of visual complexity may reflect a shift from early visual processing toward higher-order visual representations. Indeed, we observed a positive relationship between visual complexity and the coverage of microstate A, which has been previously related to activation in the temporal cortex and auditory network (Custo et al., 2017; Tarailis et al., 2023), but also to internal visualization of complex images (Milz et al., 2016). Microstate A showed both a significant positive linear and quadratic association with visual complexity, i.e., a relatively stable level at the simple visual experience level, followed by a marked increase during highly complex hallucinations. This suggests that increased recruitment of higher-order perceptual processing, maybe also language processing, contributes to complex visual hallucinations.

Besides, we observed that the explained variance of microstate C followed a U-shaped pattern with the complexity of visual hallucinations, being high at baseline (no visual experience), reaching its minimum at moderate levels of visual complexity (simple hallucinations), and increasing again at higher levels (complex hallucinations). Previous studies have linked microstate C to the salience network (Britz et al., 2010) and involved in detection of and orientation to both internal events and external stimuli. For example, increased contribution of microstate C has been associated with higher hallucination scores in adolescents with 22q11.2 syndrome (Tomescu et al., 2014), hypnosis state (Katayama et al., 2007) and patients with schizophrenia, presumably indicating aberrant salience processing (De Pieri et al., 2025; Rieger et al., 2016). From this perspective, the reduced contribution of microstate C during simple hallucination might indicate that these percepts are primarily driven by activity within sensory systems. In return, the increased microstate C during complex hallucination might suggest increased recruitment of salience-related processing as percepts get more complex and meaningful. This fits the idea that complex hallucinations arise from a variety of processes involving the frontal cortex, parietal cortex, visual systems, and thalamocortical interactions in clinical reports (Manford & Andermann, 1998). Our results might thus indicate a potential shared mechanism underlying complex hallucinations in clinical individuals and healthy individuals during MMFG.

Additionally, we found a negative linear relationship between the coverage of microstate E and visual complexity. The functional role of microstate E might correspond to a specialized aspect of salience related to interoceptive processing (Tarailis et al., 2023). Given the hypothesis that decreased interoceptive precision could lead to biased sensory sampling, disruptions in predictions and a bias towards attributing signals to external causes, which together lead to vivid and emotionally valent hallucinations (Dijkstra et al., 2024), decreased appearance of microstate E may indicate that the disruption in interoceptive processing contributes to complex visual hallucinations.

When participants reported their visual experiences, we further asked whether they experienced these events as more externally perceived perceptions or internally generated thoughts, even though they knew there were no precise sensory inputs. We observed a decrease in the explained variance for microstate C as participants increasingly believed that their experience was perception rather than thought. During MMGF, participants assumedly generated predictions and internally constructed visual representations of objects or scenes continuously, but at some points, these internally generated images crossed a ‘threshold’ and were projected externally, leading participants to mistakenly perceive them as externally generated percepts. In this context, it is interesting that microstate C has been associated with the salience network (Britz et al., 2010), which has been associated with reality monitoring (Ham et al., 2013; Oliveira et al., 2007). The decrease of microstate class C with increasing perceptual belief may thus suggest a failure in source monitoring or reality checking, thereby increasing the likelihood that participants externalize their internally generated thoughts as perceptions. This also fits with the source-monitoring accounts of hallucinations, where internally generated events are mistakenly attributed to external sources (Bentall, 1990; Ford & Hoffman, 2013; Johnson et al., 1993).

For perceptual belief during MMGF, we observed a U-shaped relationship between perceptual belief and the coverage and explained variance of microstate E, which decreased to a minimum when participants reported that their experiences were half perception and half thoughts, and increased when they had certainty about their experience (thoughts or perception). Microstate class E has previously been linked to interception, emotion, but also saliency processing (Tarailis et al., 2023). One possible interpretation is that the increased contribution of microstate E might reflect the resolution of uncertainty or imprecision in subjective states, so that participants had a clearer attribution of their experiences, regardless of whether an experience was perceived as internal or external. This resolution of uncertainty may result in reduced saliency and interoceptive conflicts. Together, these results might suggest that source monitoring during MMGF may involve two partially dissociable processes, one with determining whether an experience originated from an internal or external source, and another with the certainty or precision of that attribution.

When participants reported increased mismatch between their thoughts or imaginations and what they perceived, the contributions of microstates B and C decreased, while microstate F increased. In previous studies, these microstates have been respectively associated with visual processing (class B), salience monitoring (class C), and internally oriented mental simulation as part of the default mode network (class F, see review in Tarailis et al., 2023). In our study, the reported mismatch could arise from two perceptual possibilities: one with a real external unstructured sensory field, and another with visual hallucinations. In the first case, the increased mismatch may reflect a healthy and natural difference between freely increasing generated internal thoughts or imaginations and a consistent, uniform sensory field. In this condition, the brain naturally recruits the default mode network for spontaneous mental simulation and thoughts, as reflected in an increased appearance of microstate F and reduced engagement of visual sensory processing and salience network, as inferred from the observed decrease in appearances of microstates B and C. In the second case, when participants reported hallucinations, the mismatch was likely from the difference between participants’ predictions and different levels of perceptual hallucinations. Unlike the first case where internally generated experience could be naturally and clearly dissociated from real sensory inputs, hallucinations mistake the internally generated thoughts as external perceptions. The increased mismatch in this context may therefore indicate that the brain’s generative model has not yet converged internal predictions on a relatively stable perceptual representation, spontaneous thoughts are still active, resulting in less engagement of visual and salience-related networks and more recruitment of the default mode network. This is consistent with our phenomenological findings that when the mismatch decreased, the boundary between predictions and perception became blurred, and complex visual hallucinations gradually appeared.

In the present study, coverage of microstate D showed a negative linear relationship with active updating, i.e. the participants effort to interpret or make sense of the new perception. A similar pattern was found in previous work by Kindler et al. (2011), who reported that the duration of microstate D was significantly shorter in periods with acute hallucinations in patients with schizophrenia who, similar to our setup were able to signal the presence and absence of the hallucinatory experience. This suggests that the shortened appearance of microstate class D may facilitate the misattribution of self-generated inner speech to external sources during hallucinations. They further proposed that microstate D was involved in error-dependent updating of inferences and beliefs about the environment (Kindler et al., 2011). Following this logic, the reduced coverage of microstate D observed during active updating might reflect a diminished capacity to evaluate interpretations of the mistaken perceptual experience. Deficits in this kind of capacity have been repeatedly reported in hallucination experiences (see review in Brookwell et al., 2013). This interpretation is consistent with our phenomenological findings, in which increased active efforts to interpret new perceptual experiences contribute to increased complexity of visual hallucinations.

In the present study, we also asked about mentation before the hallucinatory experience emerged, i.e., the extent to which participants reported thinking about or imagining something. We observed that prior mentation was associated with opposite quadratic effects on the coverage and explained variance of microstates A and B: microstate A showed a negative quadratic relationship (inverted-U), whereas microstate B showed a positive quadratic relationship (U-shaped). Previous studies have found that microstate A was associated with auditory and language processing, and microstate B with visual processing (Tarailis et al., 2023). Given that people rely on both the verbal and visual modes of thoughts (Amit et al., 2013; Paivio, 1990), these findings may reflect a transition between different modes of mental representation. At moderate levels of prior mentation, participants may primarily engage in verbal thought, leading to an increased contribution of microstate A and a reduced contribution of microstate B. Then, consistent with evidence that people tend to generate visual images of what they think about verbally (Amit et al., 2017), as the intensity of mentation increases, these internally generated thoughts may become progressively elaborated into mental imagery. The increased contribution of the “visual” microstate B and the decreased contribution of the “verbal” microstate A at higher levels of prior mentation suggest a shift from verbal thought toward increasingly vivid visual representations. Together, these findings indicated that the interaction effect between verbal and visual thoughts was modulated by the intensity of participants’ ongoing mentation.

Together, the present study demonstrates that hallucinatory visual experiences during MMGF were associated with different aspects of ongoing mental experiences and were reflected by distinct EEG microstate dynamics. Some of the results indicate a potential shared mechanism underlying complex hallucinations in clinical individuals and healthy individuals during MMFG. These findings suggest that the sensory deprivation paradigm might be a valuable model for investigating the mechanisms underlying hallucinatory experiences in healthy populations and might also apply across psychosis.

## Methods

### Participants

A total of 71 healthy participants (37 males; mean age 21.4 ± 2.9 years, range 18–39 years) were recruited from the Institute of Sport Science at the University of Bern. All participants reported normal or corrected-to-normal vision and normal hearing. Exclusion criteria included a history of neurological, psychiatric, or sleep disorders, as well as the use of psychoactive or hypnotic substances, engagement in shift work. For the EEG analysis, 17 participants were excluded from further analysis due to incomplete experimental sessions (e.g., sleepiness or discomfort) or poor EEG data quality. All participants provided written informed consent prior to participation and received course credit as compensation. The study protocol was approved by the Ethics Committee of the University of Bern, and was conducted in accordance with the Declaration of Helsinki.

### Ganzfeld setting

The multimodal Ganzfeld setting is shown in Fig.1a. To create a uniform and unstructured visual field, white ping-pong balls were cut in half and shaped to fit the participants’ eye orbits comfortably. The halved ping-pong balls were then secured with transparent tape and supplemented by cotton pods to smooth the edges where necessary. Participants were seated in a comfortable chair in a darkened room. A soft, comfortable pillow was placed on their lap to support their arms and help them relax throughout the session. The lights were generated by a red LED source (Zecto Drive Max 400+ Rear, 400 lumens) and placed approximately 70cm in front of the participant’s face. Participants then listened to white noise (continuous, monotonous rain sound) delivered via over-ear headphones, which partially attenuated environmental noise. Sound intensity was individually adjusted to a subjectively comfortable level. Participants were instructed to sit in a comfortable position.

### Ganzfeld session and experience sampling

The Ganzfeld session lasted around 60 minutes. Participants were exposed to the Ganzfeld setting and instructed to sit in a relaxed manner with their eyes open. The experimental protocol was programmed using PsychoPy 2023.2.3, which controlled auditory stimulus presentation, timing, and response collection. Participants were instructed to indicate their visual experience using a mouse. Specifically, they were asked to press and hold the left mouse button while perceiving simple or abstract visual phenomena (e.g., color changes, basic shapes or patterns such as dots, lines, or flashes) that did not clearly respond to real-world objects, were difficult to describe, or lacked identifiable meaning, and to release the button when these experiences were no longer present. During the session, the protocol was interrupted by a beep at random intervals ranging from 3 to 10 minutes. During these interruptions, the audio would stop, and participants were asked to report their subjective experiences before the beep using a set of questions (Fig.1). In addition, they were instructed to click the right mouse button after they perceived complex visual phenomena which corresponded to real-world objects or environments (e.g., faces, animals, or plants), or when they experienced being in a different space or situation, such as a street, forest, or another room, even being inside a dream-like environment. After clicking the right mouse button, a beep sounded, the ongoing protocol was interrupted, and participants were asked to report their experience before the beep using the same set of questions. To facilitate longer and more vivid complex visual experiences, participants were encouraged to click the right button after the experience had persisted for at least 10 seconds.

To quantify participants’ subjective experiences during the Ganzfeld session, we asked a set of questions at random intervals (Supplementary Note 1). The questions were inspired by the Prediction-Related Experiences Questionnaire (O’Brien et al., 2024), which was developed to assess subjective experiences of prediction in everyday life. Participants were asked about: (1) visual complexity: What sensory experience did you have before the beep; (2) perceptual belief: Which was more prevalent in your experience, thoughts, or perceptions (rated on a scale from 0 = completely thoughts to 4 = completely perception); (3) Prior mentation: before the experience occurred/beep, to what extent were you thinking about or imagining something (Rated on a scale from 0 = not at all to 4 = very much); (4) Perception prediction mismatch: How closely did your thoughts or imagination match what you perceived (Rated on a scale from 0 = not at all to 4 = very much). For the analyses, these ratings were reverse-coded so that higher values reflected greater mismatch between mentation and perception (0 = no mismatch; 4 = very high mismatch) ; (5) Active Updates: How strongly did your mind try to interpret or make sense of the new perception? (Rated on a scale from 0 = not at all to 4 = very much).

### Experimental procedures

Prior to the laboratory session, Participants were reminded to have a normal wake-sleep rhythm as usual, no drugs and no alcohol 24 hours before the experiment. Upon arrival at the laboratory, participants were introduced to the experiment procedure, gave their informed consent and received and gave relevant documentation (Pseudonymous code and demographic data). They subsequently completed measures of sleepiness (Johns, 1991) and handedness. Participants who reported high levels of sleepiness (sleepiness score≥ 8) were advised to reschedule the session.

Participants were then prepared for the EEG recording. During this period, the procedure and tasks of the Ganzfeld session were explained to participants in detail. After the EEG setup was completed, participants sat in a chair in a comfortable position. Before starting the protocol, an EEG eyemovement calibration was recorded, during which participants performed a series of eye movements, including blinking, vertical eye movements (up and down), and horizontal eye movements (left and right). Next, a 2-minute resting-state EEG was recorded, consisting of 20 seconds with eyes open and 40 seconds with eyes closed, repeated twice. Then, participants were fitted with halved ping-pong balls placed over their eyes and were instructed to keep their eyes open and blink normally. Continuous white noise was presented via the headphones, with the volume adjusted to a comfortable level. Participants held a mouse in their right hand.

To familiarize the participants with the experimental protocol and to control for EEG activity associated with motor responses (button presses during a simple visual experience), a control button-click task was performed. Participants were instructed to simulate the button-press task associated with simple visual experiences by pressing and holding the left mouse button for 10 seconds, followed by a 10-second rest period. This sequence was repeated for a total duration of 2 minutes. Data from this task were not included in the analyses in the present study. Then the Ganzfeld session began, during which participants’ subjective experiences were sampled at random intervals using a set of questions. Participants were instructed to keep their eyes open, blink normally, relax, and allow their thoughts and imagery to flow freely during the Ganzfeld session. They were also informed that they could terminate the session at any time if they experienced discomfort.

### EEG Recording

EEG signals were recorded from 64 scalp electrodes using a sponge-based R-Net system (Brain Products GmbH, Gilching, Germany) according to the 10-20 system. Different cap sizes were applied depending on the participant’s head circumference. All electrodes were online referenced to the reference electrode on the FCz and grounded to the Fpz. The recording setup employed two LiveAmp32 from BrainProducts (Brain Products GmbH, Gilching, Germany). The recording was conducted using BrainVision Recorder software (version 1.27) at a sampling rate of 250 Hz. The electrode impedance was kept below 50 kΩ.

### Data analysis

#### Experience sampling analysis

Participants’ reported visual experiences from experience sampling trials were categorized by the authors according to their spatial characteristics, following the framework of Lloyd et al. (2012). These experiences includes a continuum of visual perceptions, from abstract simple perceptions: 1) simple visual changes, e.g., color changes, flash; 2) Simple shapes or patterns e.g., lines, dots, cloudy and 3) Combine 1 and 2, e.g., a sequence of dots moving like a river; to external object related and immersive complex perceptions: 4) objects, e.g., face, tree, ghost and 5) environment, e.g., sea, garden or dream liked experiences. Trials in which participants reported no sensory experience were assigned a baseline score of 0 (Fig. 2).

### EEG Preprocessing

EEG data preprocessing was performed using BrainVision Analyzer 2.2 (BrainProducts GmbH, Gilching, Germany) and MATLAB R2023b (Mathworks Inc. Natick, MA, USA). Bad channels were identified by visual inspection and interpolated using spherical splines. The continuous EEG data were first segmented to include only the Ganzfeld periods, excluding intervals corresponding to experience-sampling reports. To make the subsequent EEG microstate analysis filtering more flexible, individual spatial filters was made to remove artifacts. To design these filters, the data were re-referenced to the average reference and band-pass filtered between 1.5 and 30 Hz. Independent component analysis (ICA) was then performed to identify components corresponding to ocular, muscular, and electrocardiographic artifacts, and a spatial filter that removed these components was build. Then, this spatial filter was applied to the initially segmented data. EEG segments corresponding to the 30 s preceding each beep probe were extracted for further analysis. The remaining physiological or technical artifacts in this data were removed by visual inspection.

### EEG Microstate analysis

The thus selected data were re-referenced to the average reference and filtered between 2 and 20 Hz. EEG microstate analysis was performed using MICROSTATELAB, an EEGLAB toolbox for resting-state microstate analysis (Nagabhushan Kalburgi et al., 2023). First, the Global Field Power (GFP) was computed for each sample over time. Topographies at the local maxima of GFP were clustered for each trial, using a fixed range of cluster numbers (4–7) and a modified k-means algorithm. The polarity of the topographies was ignored. Subsequently, mean microstate topographies were identified across trials and participants for each k. We then sorted these sets of mean microstate topographies based on published template maps (Custo et al., 2017) and saved them as template maps. Because we had an explicit hypothesis about microstates B, C, and D and visual experience during MMGF. Only the seven-class solution included all the microstate class maps (microstates A, B, C, D, E, F, G) we hypothesized. Therefore, after cluster analysis, we selected seven microstates to capture the effects of experience changes during MMGF. Temporal features of these identified microstates were then extracted, including the average coverage of the EEG signal in percent for each microstate class (Coverage), and the explained variance for each microstate class (ExpVar).

### Meta-microstate Analysis

Meta-microstate analysis on seven microstate classes were conducted with MATLAB applications MSTemplateEditor and MSTemplateExplorer (https://github.com/ThomasKoenigBern/MS-Template-Explorer) (Koenig et al., 2023). We assessed the topographical similarities between our microstate template maps and those from other studies in the database, generating a similarity matrix with the MSTemplateExplorer app. Then we selected empirical findings from the database associated with microstate maps that exhibited topographic similarity with our template maps and were conceptually connected to our hypotheses.

### Statistical analysis

To assess potential associations between distinct microstate networks and experience-sampling ratings, the ratings were decomposed into linear and quadratic (poly, degree = 2) terms. Then we performed separate linear mixed models for the dependent variables of coverage and explained variance. The linear and quadratic polynomial terms and microstate class were used as fixed factors. A by-subject random intercept was added to represent between-person variability and account for the unbalanced data structure (Bates et al., 2015).

For the phenomenological data, we examined the associations between visual complexity and perceptual belief, prior mentation, perception-prediction mismatch and active updating. We fitted separate linear mixed-effects models with collected ratings as the dependent variables and visual complexity terms as the predictor. The linear and quadratic polynomial terms were used as fixed factors, with subject as a random factor. We analyzed the data using R version 4.6.0, fitting all models with the package lme4 (Bates et al., 2015). ANOVA tests were performed to assess the significance of each linear mixed-effects model. Post hoc comparisons were estimated using ‘emtrends’ and corrected for multiple comparisons using the false discovery rate (FDR).

## Author contributions

X.W., D.E., and T.K. designed the research; X.W. and Y.P. collected the data; X.W. and T.K. analyzed data; X.W. wrote the original draft; E.P., D.E., and T.K. reviewed and edited the manuscript.

## Declaration of generative AI and AI-assisted technologies in the writing process

During the preparation of this work, the author(s) used Grammarly and ChatGPT to check and improve grammar and sentence structure. After using this tool, the author(s) reviewed and edited the content as needed and take full responsibility for the content of the publication.

