## Supplemental file for "Multimodal Ganzfeld-induced visual experiences are associated with accompanying mental experiences and distinct EEG microstate dynamics"

### **Supplementary Note 1.** Random interval experience sampling questions (RIES)

1. What sensory experience did you have before the beep?

(0 - none, 1 - simple perception: color changes; 2 - Simple shapes or patterns: lines, dots, cloudy; 3 - 1 and 2, or moving like a river....; 4 - objects: face, tree, ghost; environment; 5 - dream liked experiences: object and environment) This part was rated by researchers.

(only ask the detailed questions when the experience ≥4)

Please rate the following statements on a 5-point scale (0 = not at all, 1 = just a little, 2 = moderately, 3 = pretty much, 4 = very much)

2. Which was more prevalent in your experience, thoughts, or perceptions? (0 is completely thoughts, 4 = completely perception)

3. Before the experience occurred/beep, to what extent were you thinking about or imagining something?

4. How closely did your thoughts or imagination match what you perceived?

5. How strongly did your mind try to interpret or make sense of the new perception?

### Microstate template maps for solutions ranging from 4 to 7 classes

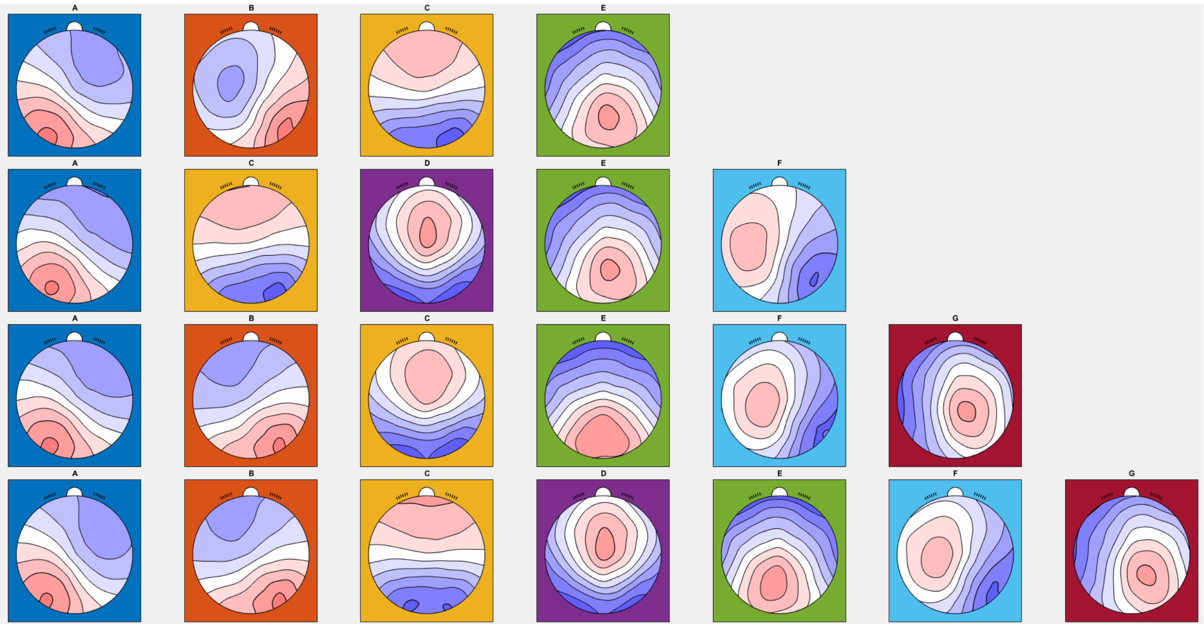


**Figure S1. Microstate template maps for solutions ranging from 4 to 7 classes.** The 4-class solution identified microstate classes A, B, C, and E. The 5-class solution identified microstate classes A, C, D, E, and F. The 6-class solution identified microstate classes A, B, C, E, F, and G. The 7-class solution identified microstate classes A, B, C, D, E, F, G.

| | A (4) | B (4) | C (4) | E (4) | A (5) | C (5) | D (5) | E (5) | F (5) | A (6) | B (6) |
|---|---|---|---|---|---|---|---|---|---|---|---|
| A (4) | **100.0000** | 0.1302 | 53.6535 | 22.0662 | **95.9664** | 46.8444 | 52.2451 | 14.8247 | 7.5780 | **98.6935** | 20.3661 |
| B (4) | 0.1302 | **100.0000** | 39.8993 | 20.2109 | 0.3422 | 42.4177 | 30.7677 | 23.2599 | 93.1191 | 0.1715 | **68.5933** |
| C (4) | 53.6535 | 39.8993 | **100.0000** | 60.8173 | 54.2258 | **98.7349** | 72.0424 | 55.0796 | 19.8506 | 55.2783 | 87.6811 |
| E (4) | 22.0662 | 20.2109 | 60.8173 | **100.0000** | 35.8383 | 70.6549 | 10.9842 | **99.1019** | 15.5797 | 30.1699 | 67.4165 |
| A (5) | **95.9664** | 0.3422 | 54.2258 | 35.8383 | **100.0000** | 50.2431 | 38.4773 | 27.1064 | 7.3527 | **99.2115** | 22.9948 |
| C (5) | 46.8444 | 42.4177 | **98.7349** | 70.6549 | 50.2431 | **100.0000** | 61.9533 | 65.6623 | 23.6449 | 49.9849 | 91.8633 |
| D (5) | 52.2451 | 30.7677 | 72.0424 | 10.9842 | 38.4773 | 61.9533 | **100.0000** | 7.6496 | 10.0883 | 45.2190 | 50.5235 |
| E (5) | 14.8247 | 23.2599 | 55.0796 | **99.1019** | 27.1064 | 65.6623 | 7.6496 | **100.0000** | 20.1964 | 21.9214 | 66.5957 |
| F (5) | 7.5780 | 93.1191 | 19.8506 | 15.5797 | 7.3527 | 23.6449 | 10.0883 | 20.1964 | **100.0000** | 7.1299 | 50.8600 |
| A (6) | **98.6935** | 0.1715 | 55.2783 | 30.1699 | **99.2115** | 49.9849 | 45.2190 | 21.9214 | 7.1299 | **100.0000** | 22.6728 |
| B (6) | 20.3661 | **68.5933** | 87.6811 | 67.4165 | 22.9948 | 91.8633 | 50.5235 | 66.5957 | 50.8600 | 22.6728 | **100.0000** |
| C (6) | 59.3018 | 34.9673 | **89.6685** | 29.0306 | 50.4384 | **82.0859** | 94.7015 | 23.7658 | 13.2870 | 55.4386 | 68.6415 |
| E (6) | 51.4926 | 12.0995 | 76.1360 | **90.5986** | 65.8255 | 81.6297 | 26.7245 | **84.4566** | 4.8819 | 60.4718 | 65.3058 |
| F (6) | 8.5466 | 86.0275 | 11.8274 | 0.7994 | 13.0542 | 12.0719 | 17.8151 | 1.7902 | **86.3903** | 10.6368 | 32.0706 |
| G (6) | 8.2642 | 22.9184 | 5.8006 | 50.3318 | 1.5903 | 12.0116 | 6.1130 | 59.5558 | 37.6850 | 3.7345 | 25.5361 |
| A (7) | **97.5638** | 3.6515 | 38.5818 | 15.3545 | **94.3687** | 32.5973 | 38.8507 | 9.2142 | 17.6870 | **96.4925** | 9.8869 |
| B (7) | 22.5019 | **67.8872** | 89.5237 | 65.3727 | 24.6454 | 92.9632 | 54.2402 | 64.0915 | 49.0679 | 24.5817 | **99.8510** |
| C (7) | 66.3924 | 24.3568 | **96.9140** | 66.1528 | 69.8210 | **95.9404** | 64.1309 | 58.8113 | 9.0990 | 69.6959 | 77.6840 |
| D (7) | 52.4210 | 25.4708 | 64.8131 | 6.6867 | 37.1312 | 54.1359 | **99.2824** | 4.0128 | 6.8944 | 44.4031 | 42.2253 |
| E (7) | 49.2545 | 0.9300 | 51.0559 | **83.2417** | 67.9822 | 56.7234 | 10.0094 | **76.8546** | 0.0015 | 60.2768 | 38.5218 |
| F (7) | 4.7882 | 93.4488 | 19.4855 | 4.5158 | 7.3953 | 20.3754 | 21.6821 | 6.4742 | **91.9350** | 5.9003 | 43.7520 |
| G (7) | 0.3150 | 39.3985 | 25.3915 | 71.9662 | 0.7290 | 35.4716 | 0.1073 | 79.9383 | 48.2499 | 0.0848 | 52.7254 |

| | C (6) | E (6) | F (6) | G (6) | A (7) | B (7) | C (7) | D (7) | E (7) | F (7) | G (7) |
|---|---|---|---|---|---|---|---|---|---|---|---|
| A (4) | 59.3018 | 51.4926 | 8.5466 | 8.2642 | **97.5638** | 22.5019 | 66.3924 | 52.4210 | 49.2545 | 4.7882 | 0.3150 |
| B (4) | 34.9673 | 12.0995 | 86.0275 | 22.9184 | 3.6515 | **67.8872** | 24.3568 | 25.4708 | 0.9300 | 93.4488 | 39.3985 |
| C (4) | **89.6685** | 76.1360 | 11.8274 | 5.8006 | 38.5818 | 89.5237 | **96.9140** | 64.8131 | 51.0559 | 19.4855 | 25.3915 |
| E (4) | 29.0306 | **90.5986** | 0.7994 | 50.3318 | 15.3545 | 65.3727 | 66.1528 | 6.6867 | **83.2417** | 4.5158 | 71.9662 |
| A (5) | 50.4384 | 65.8255 | 13.0542 | 1.5903 | **94.3687** | 24.6454 | 69.8210 | 37.1312 | 67.9822 | 7.3953 | 0.7290 |
| C (5) | **82.0859** | 81.6297 | 12.0719 | 12.0116 | 32.5973 | 92.9632 | **95.9404** | 54.1359 | 56.7234 | 20.3754 | 35.4716 |
| D (5) | 94.7015 | 26.7245 | 17.8151 | 6.1130 | 38.8507 | 54.2402 | 64.1309 | **99.2824** | 10.0094 | 21.6821 | 0.1073 |
| E (5) | 23.7658 | **84.4566** | 1.7902 | 59.5558 | 9.2142 | 64.0915 | 58.8113 | 4.0128 | **76.8546** | 6.4742 | 79.9383 |
| F (5) | 13.2870 | 4.8819 | **86.3903** | 37.6850 | 17.6870 | 49.0679 | 9.0990 | 6.8944 | 0.0015 | **91.9350** | 48.2499 |
| A (6) | 55.4386 | 60.4718 | 10.6368 | 3.7345 | **96.4925** | 24.5817 | 69.6959 | 44.4031 | 60.2768 | 5.9003 | 0.0848 |
| B (6) | 68.6415 | 65.3058 | 32.0706 | 25.5361 | 9.8869 | **99.8510** | 77.6840 | 42.2253 | 38.5218 | 43.7520 | 52.7254 |
| C (6) | **100.0000** | 48.4861 | 14.7019 | 0.3184 | 44.1154 | 72.0308 | **83.7148** | 90.7833 | 26.0964 | 20.4863 | 5.1844 |
| E (6) | 48.4861 | **100.0000** | 0.0608 | 21.1209 | 42.3767 | 65.0092 | 85.8393 | 21.1488 | **92.4549** | 0.9576 | 42.3172 |
| F (6) | 14.7019 | 0.0608 | **100.0000** | 8.3818 | 18.6331 | 31.8267 | 3.0589 | 15.5208 | 7.9635 | **98.3947** | 14.5853 |
| G (6) | 0.3184 | 21.1209 | 8.3818 | **100.0000** | 12.6960 | 22.3456 | 4.7691 | 10.6689 | 19.4828 | 13.2960 | **91.9917** |
| A (7) | 44.1154 | 42.3767 | 18.6331 | 12.6960 | **100.0000** | 11.4085 | 52.3535 | 39.9778 | 45.5223 | 13.3662 | 2.3203 |
| B (7) | 72.0308 | 65.0092 | 31.8267 | 22.3456 | 11.4085 | **100.0000** | 79.4409 | 45.9078 | 37.9463 | 43.3343 | 48.9893 |
| C (7) | **83.7148** | 85.8393 | 3.0589 | 4.7691 | 52.3535 | 79.4409 | **100.0000** | 57.5770 | 65.6300 | 7.8368 | 22.5381 |
| D (7) | 90.7833 | 21.1488 | 15.5208 | 10.6689 | 39.9778 | 45.9078 | 57.5770 | **100.0000** | 6.9822 | 18.4124 | 0.2525 |
| E (7) | 26.0964 | **92.4549** | 7.9635 | 19.4828 | 45.5223 | 37.9463 | 65.6300 | 6.9822 | **100.0000** | 2.5875 | 33.9183 |
| F (7) | 20.4863 | 0.9576 | **98.3947** | 13.2960 | 13.3662 | 43.3343 | 7.8368 | 18.4124 | 2.5875 | **100.0000** | 22.5804 |
| G (7) | 5.1844 | 42.3172 | 14.5853 | **91.9917** | 2.3203 | 48.9893 | 22.5381 | 0.2525 | 33.9183 | 22.5804 | **100.0000** |

**Figure S2. The similarity matrix obtained from 4-7 solutions of template maps.** Each column and row represented one template map of one class solution. The value in the cell represented the topographical similarities between two maps at that row and column. The color bar indicates that darker shades of green in the matrix cells correspond to higher spatial similarity between pairs of maps.

### Meta-microstate Analysis Results

#### Supplementary Table 1. Summary of empirical findings for microstate A

| Study | Comparison / localization | findings | Eyes | Similarity |
|---|---|---|---|---|
| (Custo et al., 2017) | Left temporal cortex (BA41/PAC, BA22/Wernicke); insula; lingual gyrus (BA19) | ↑ Current density | EC | 99.0% |
| (Milz et al., 2016) | Object/spatial visualization vs verbalization | ↑ Coverage | EC | 98.2% |
| (Damborská, Piguet, et al., 2019) | Euthymic bipolar disorder vs controls | ↑ Coverage | EC | 97.8% |
| (Zanesco et al., 2020) | Negatively to Alertness reaction time | ↑ Duration | EC+EO | 97.4% |
| (Giordano et al., 2018) | Increasing avolition–apathy, anhedonia, avolition, and asociality in patient with schizophrenia | ↑ Coverage | EC | 96.8% |
| (Tomescu et al., 2022) | Social imitation vs control; planning/verbal thoughts; high vs low control-task performance | ↑ Duration; ↑ occurrence | EC | 96.5% |
| (Damborská, Tomescu, et al., 2019) | Depressive symptomatology (MADRS) | ↑ Occurrence | EC | 96.1% |
| (Smailovic et al., 2019) | cognitive decline | ↑Duration; ↑ Coverage | EC | 95.6% |
| (Hanoglu et al., 2022) | Alzheimer's disease vs controls | ↑ Coverage; ↑ occurrence | EC | 94.8% |
| (Lehmann et al., 2005) | Schizophrenia vs controls | ↑ Occurrence; ↑ Coverage | EC | 93.6 % |
| (Musaeus et al., 2020) | Mild cognitive impairment and Alzheimer's dementia vs healthy controls | ↑ Duration; ↑ Coverage; ↑ occurrence | EC | 91.5% |
| (Britz et al., 2010) | Bilateral superior/middle temporal gyri and left middle frontal gyrus | ↓ BOLD | EC | 90.4% |

* ↑, higher/increased; ↓, lower/decreased; ARSQ, Amsterdam Resting-State Questionnaire; BA, Brodmann area; BOLD, blood-oxygen-level-dependent signal; EC, eyes closed; EO, eyes opened.

#### Supplementary Table 2. Summary of empirical findings for microstate B

| Study | Comparison / localization | findings | Eyes | Similarity |
|---|---|---|---|---|
| (Custo et al., 2017) | Bilateral occipital cortices (cuneus; BA17/BA18); right insular cortex extending to right claustrum and right frontal eye field (BA8) | ↑ Current density | EC | 92.8% |
| (Schiller et al., 2021) | Intoxicated vs sober | ↑ Contribution | EC | 90.2% |
| (Qin et al., 2022) | High vs low intensity of depressive symptoms | ↑ Duration; ↑ occurrence; ↑ coverage | EC | 89.1% |
| (Lehmann et al., 2005) | Schizophrenia vs controls | ↑ Occurrence | EC | 86.9% |

* ↑, higher/increased; ↓, lower/decreased; ARSQ, Amsterdam Resting-State Questionnaire; BA, Brodmann area; EC, eyes closed; EO, eyes opened.

#### Supplementary Table 3. Summary of empirical findings for microstate C

| Study | Comparison / localization | findings | Eyes | Similarity |
|---|---|---|---|---|
| (Lehmann et al., 2005) | Patient with Schizophrenia vs controls | ↑ Occurrence | EC | 99.5% |
| (Spring et al., 2018) | Endurance exercise vs baseline | ↑ Duration; ↑ contribution | EC | 98.9% |
| (Custo et al., 2017) | Precuneus/posterior cingulate cortex (PCC) and left angular gyrus | ↑ Current density | EC | 98.2% |
| (Tarailis et al., 2021) | Higher ARSQ Comfort score | ↓ Occurrence | EC | 97.8% |
| (Britz et al., 2010) | Bilateral ACC, medial cingulate gyrus, left inferior frontal gyrus, left claustrum, right frontal gyrus, and right amygdala | ↑ BOLD | EC | 97.8% |
| (Zanesco et al., 2021) | Mind-wandering vs on-task | ↑ GEV; ↑ GFP; ↑ contribution | EC | 94.7% |
| (Müller et al., 2005) | Before vs after perceptual flip | ↑ Contribution before 750 ms; ↓ contribution after 300 ms before perceptual flip | EO | 94.6% |
| (Spring et al., 2017) | 30-min endurance exercise vs pre-exercise | ↑ GEV; ↑ duration; ↑ contribution | EC | 94.3% |
| (Milz et al., 2016) | Resting vs object visualization; resting vs verbalization | ↑ Occurrence; ↑ contribution | EC | 93.8% |
| (Croce et al., 2022) | Isometric contraction task vs rest | ↑ Duration; ↑ occurrence | EO | 91.3% |
| (Tomescu et al., 2022) | Social imitation vs control task in subjects with higher oxytocin | ↑ Duration | EC | 91.0% |
| (Murphy et al., 2020) | First-episode psychosis vs controls | ↓ Contribution | EC | 90.9% |
| (Zappasodi et al., 2019) | Spatial relation/visualization vs induction; spatial relation task vs induction | ↓ Contribution; ↓ duration | EO | 89.5% |
| (Toplutaş et al., 2024) | Disorders of consciousness vs healthy controls | ↑ Duration; ↑ contribution | EO | 89.3% |
| (Giordano et al., 2018) | Schizophrenia vs controls | ↑ Contribution; ↑ duration | EC | 85.8% |
| (Schiller et al., 2021) | Intoxicated vs sober | ↓ Contribution; ↓ duration | EC | 85.2% |
| (Denzer et al., 2024) | Bizarre vs normal VR environment (trend) | ↓ Contribution | EO | 84.8% |

* ↑, higher/increased; ↓, lower/decreased; ACC, anterior cingulate cortex; GEV, global explained variance; GFP, global field power; BOLD, blood-oxygen-level-dependent signal; VR, virtual reality; EC, eyes closed; EO, eyes opened.

**Supplementary Table 4. Summary of empirical findings for microstate D**

| Study | Comparison / localization | findings | Eyes | Similarity |
|---|---|---|---|---|
| (Tarailis et al., 2021) | Higher ARSQ score Self | ↓ Duration | EC | 98.4% |
| (Spring et al., 2018) | endurance excercise against baseline | ↑ Occurrence | EC | 97.5% |
| (Smailovic et al., 2019) | Cognitive Decline | ↓ Occurrence; ↓ Contribution | EC | 94.4% |
| (Croce et al., 2018) | TMS post against pre stimulation on right side IPS | ↓ Contribution; ↓ Duration; ↓ Occurrence | EO | 94.2% |
| (Hu et al., 2023) | Post against pre visual emotion-evoking task; high against low emotional valence | Post against pre visual emotion-evoking task: ↑ Contribution; high against low emotional valence: ↓ Occurrence | EO | 93.9% |

| | | | | |
|---|---|---|---|---|
| (Hanoglu et al., 2022) | Alzheimer's and parkinson's against controls; parkinson's against controls | Alzheimer's and parkinson's against controls: ↓ Duration; parkinson's against controls: ↓ Contribution | EC | 92.7% |
| (Zanesco et al., 2020) | More extraversion, agreeableness, conscientiousness | ↑ GEV, ↑ Occurrence | EC+EO | 89.4% |
| (Lehmann et al., 2005) | Patients with Schizophrenia against Controls | ↓ Duration | EC | 88.9% |
| (Bréchet et al., 2020) | Slowwave sleep vs wakeful resting; Dream experience vs no experience in slowwave sleep | ↑ GEV | EC | 86.8% |
| (Britz et al., 2010) | Right superior and inferior parietal lobules; Right superior and middle frontal gyrus | ↓ BOLD | EC | 84.3% |
| (Zappasodi et al., 2019) | Spatial relationship task (fluid intelligence) against control items or visualization and induction | ↑ Contribution | EO | 83.7% |
| (Custo et al., 2017) | Right inferior parietal lobe (BA40); Right middle and superior frontal gyri; Right insula (BA13) | ↑ Current density | EC | 83.4% |

* ↑, higher/increased; ↓, lower/decreased; ARSQ, Amsterdam Resting-State Questionnaire; BOLD, blood-oxygen-level-dependent signal; GEV, global explained variance; GFP, global field power; TMS, transcranial magnetic stimulation; IPS, intraparietal sulcus; EC, eyes closed; EO, eyes opened.

**Supplementary Table 5. Summary of empirical findings for microstate E**

| Study | Comparison / localization | findings | Eyes | Similarity |
|---|---|---|---|---|
| (Custo et al., 2017) | Dorsal anterior cingulate cortex (BA32) extending to superior frontal gyrus; bilateral middle frontal gyri; bilateral insulae | ↑ Current density | EC | 95.7% |
| (Hu et al., 2022) | After vs before watching an emotional video; higher valence rating after watching | Post vs pre: ↓ contribution, duration, occurrence; higher valence: ↑ contribution and occurrence | EO | 94.0% |
| (Hu et al., 2023) | Post- vs pre-visual emotion-evoking task; high vs low emotional arousal | Post vs pre: ↓ contribution, duration, occurrence; high arousal: ↑ contribution and occurrence | EO | 94.0% |
| (Denzer et al., 2024) | Bizarre vs normal VR environment | ↑ Contribution; ↑ GFP (trend) | EO | 90.2% |
| (Giordano et al., 2018) | Schizophrenia vs controls | ↑ Contribution; ↑ duration | EC | 89.6% |
| (Tarailis et al., 2021) | Higher ARSQ Somatic Awareness score | ↓ Contribution | EC | 89.5% |
| (Zappasodi et al., 2019) | Spatial relation/visualization vs induction | ↓ Contribution; ↓ duration | EO | 87.6% |
| (Tomescu et al., 2022) | High vs low control-task performance; planning and verbal thoughts | ↑ Duration with high performance; ↑ occurrence negatively associated with planning/verbal thoughts | EC | 87.4% |

| | | | | |
|---|---|---|---|---|
| (Zanesco et al., 2020) | Conscientiousness and nervous mood | ↑ GEV with higher conscientiousness and lower nervous mood; ↑ occurrence with higher conscientiousness | EC+EO | 87.2% |
| (Zanesco et al., 2021) | Mind-wandering vs on-task | ↓ GEV; ↓ occurrence; ↓ contribution | EC | 87.2% |

* ↑, higher/increased; ↓, lower/decreased; ARSQ, Amsterdam Resting-State Questionnaire; BA, Brodmann area; GEV, global explained variance; GFP, global field power; VR, virtual reality; EC, eyes closed; EO, eyes opened.

**Supplementary Table 6. Summary of empirical findings for microstate F**

| Study | Comparison / localization | findings | Eyes | Similarity |
|---|---|---|---|---|
| (Qin et al., 2022) | High vs low intensity of depressive symptoms | ↓ Occurrence; ↓ coverage | EC | 90.6% |
| (Tarailis et al., 2021) | Higher ARSQ Comfort score | ↑ Duration | EC | 90.5% |
| (Tomescu et al., 2022) | Planning and verbal thoughts | ↑ Occurrence (positive association) | EC | 84.3% |
| (Custo et al., 2017) | Left middle frontal gyrus/frontal eye field (BA8), dorsal anterior cingulate, cuneus extending to PCC, and thalamus | ↑ Current density | EC | 81.1% |
| (Diezig et al., 2022) | Hypnagogic state vs wake; right-lateralized temporal, occipital, fusiform and cerebellar activity | ↓ Contribution; ↓ duration; ↓ occurrence; ↑ current density | EC | 75.0% |
| (Hu et al., 2023) | Post- vs pre-visual emotion-evoking task | ↑ Contribution | EO | 74.8% |
| (Britz et al., 2010) | Bilateral inferior occipital gyri and cuneus, left lingual gyrus, middle occipital gyrus | ↓ BOLD | EC | 73.9% |

* ↑, higher/increased; ↓, lower/decreased; ARSQ, Amsterdam Resting-State Questionnaire; BOLD, blood-oxygen-level-dependent signal; GEV, global explained variance; GFP, global field power; PCC, posterior cingulate cortex; EC, eyes closed; EO, eyes opened.

**Supplementary Table 7. Summary of empirical findings for microstate G**

| Study | Comparison / localization | findings | Eyes | Similarity |
|---|---|---|---|---|
| (Croce et al., 2022) | Isometric contraction task vs rest | ↑ Duration; ↑ occurrence | EO | 90.7% |
| (Zappasodi et al., 2019) | Spatial relationship/visualization vs control or induction; induction vs visualization/control | ↑ Contribution for spatial relationship/visualization; ↓ duration and occurrence for induction contrasts | EO | 84.1% |
| (Custo et al., 2017) | Right inferior parietal lobe extending to superior temporal gyrus; cerebellum | ↑ Current density | EC | 83.7% |

* ↑, higher/increased; ↓, lower/decreased; EC, eyes closed; EO, eyes opened
